# FINET: Fast Inferring NETwork

**DOI:** 10.1101/733683

**Authors:** Anyou Wang, Rong Hai

**Author notes:** Correspondence: A Wang, R Hai.

## Abstract

Numerous software have been developed to infer the gene regulatory network, a long-standing key topic in biology and computational biology. Yet the slowness and inaccuracy inherited in current software hamper their applications to the increasing massive data. Here, we develop a software, FINET (<u>F</u>ast Inferring <u>NET</u>work), to infer a network with high accuracy and rapidity. The high accuracy results from integrating algorithms with stability-selection, elastic-net, and parameter optimization. Tested by a known biological network, FINET infers interactions with more than 94% precision (true positives/total true callings). The high speed comes from partnering parallel computations implemented with Julia, a new compiled language that runs much faster than existing languages used in the current software, such as R, Python, and MATLAB. Regardless of FINET’s implementations with Julia, users without any background in the language or computer science can easily operate it, with only a user-friendly single command line. In addition, FINET can infer other networks such as chemical networks and social networks. Overall, FINET provides a confident way to efficiently and accurately infer any type of network for any scale of data.

Availability and implementation available in github https://github.com/anyouwang/finet.git

## 1. Introduction

All biological phenotypes result from a certain degree of gene regulation, and understanding gene regulations remains a crucially fundamental topic in the biology. Conventionally, manipulating gene mutations such as knockout and knockdown helps to infer the gene regulations. However, these approaches suffer several challenges such as transcript compensatory and side effects ^1^. Gene mutation approaches also assume that the genome becomes stable after mutations. The genome, however, varies dramatically with even a single gene mutation, which alters gene expressions of thousand genes as shown in RNA sequencing data. As a result, there is no way to fully comprehend the complete regulatory interactions of any single gene.

Computational biology and bioinformatics have attempted to infer gene regulatory networks from gene expression data, and have established software and tools to execute their works ^2–7^. However, the efficiency of current software suffers from high noise and lagging, and they usually generate overly complicated network interactions—mostly false positives ^2^. Therefore, these results actually provide more questions than answers to true biology regulatory interactions. With the software FINET, we are able to quickly and accurately reveal true gene interactions and refresh gene interaction pictures.

## 2. Theory and algorithms

Theoretically, FINET is primarily based on elastic-net theory and stability selections. The elastic-net is an extension of LASSO^8^ (least absolute shrinkage and selection operator), a penalized regression method for shrinkage and variable selection by minimizing:

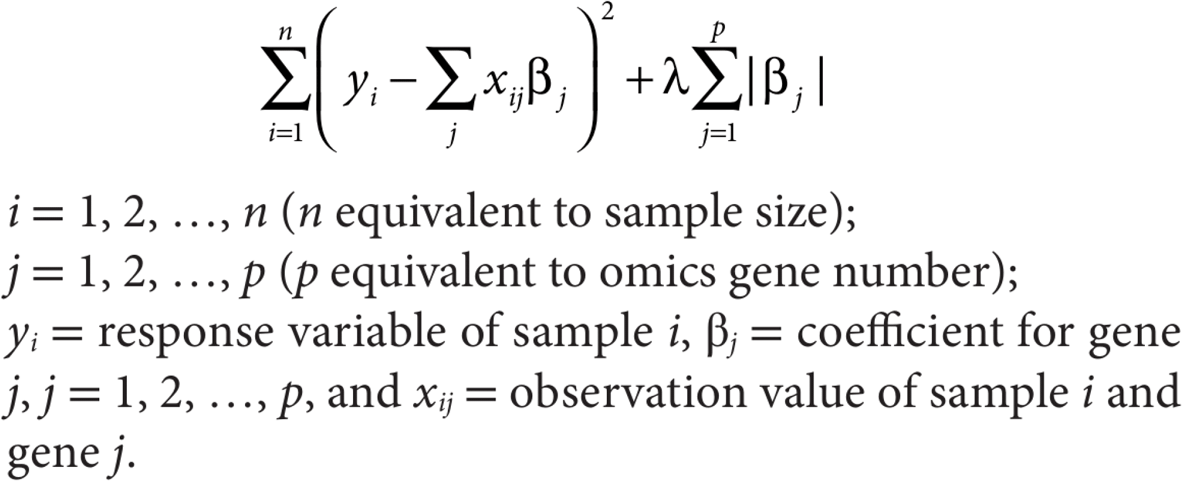

Lasso tends to ignore the variables in a correlated group. To avoid this, the elastic-net adds an additional quadratic part 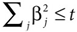 to the penalization to include the correlated genes.

Elastic-net and lasso are arguably the best methods for shrinkage and variable selection, and k-fold cross-validations have been implemented in current software like GMLNet ^9^. However, these validations include too many variables and these selected variables offer results of coefficients without any priority of trueness. It is then difficult to estimate the stability of these variable selections.

To improve the accuracy of variable selection, stability selection comes into play ^10^. The general idea of stability selection is to add a resampling step into an existing model selection to make it stable and increase accuracy. For example, during elastic-net selection, the total samples are randomly partitioned into two subgroups, and each subgroup is subjected to an elastic-net model selection. If a variable was simultaneously selected at the two groups, the selected variable would be likely true ^10^.

The FINET’s algorithm of each resampling step is to bootstrap randomly split samples into m subgroups (m >=2) without replacement. In each subgroup, a complete model of elastic-net is run to select variables (regulators in biology) interacting with a target (a target gene in biology). Such resampling step iterates *n* times. The frequency of each regulator selected during iterations is counted as frequency score, total selected times in n*m trials = total hits/n*m, and it is used to rank regulator priority of confidences (frequency levels) and confidence strength in true positive selection. The maximum frequency score is 1 (highest confidence), and a variable with a frequency of 1 for a given target means that it was always selected in *m*n* trials and it is likely a true positive regulator for this target. When m increases (e.g. m=8), in which a regulator simultaneously targets its target at m sub-groups in n bootstrap resampling, type I error goes down dramatically.

## 3. Parameter optimization

We have optimized FINET parameters for most common users and was set as default values in FINET. Here, we only highlighted parameter optimization of the frequency score cutoff and resampling in *m* groups.

### 3.1. Frequency score cutoff

To systematically optimize the frequency score cutoff for FINET, we run FINET to select regulators controlling each target in a well-known matrix used by dream5 network challenge (network1 matrix) ^2^, which includes variable matrix and golden standard true positives.

From the theory above, we learned that high frequency cutoff ensures the accuracy of variable selection. The optimal cutoff, however, remains unknown. To optimize the frequency cutoff, we first computed the AUC (Area Under The Curve) of ROC (receiver operating characteristic curve) at an array of frequency from 0.1 to 1. The golden standard at network1 was treated as known interactions, and the total true positives produced by FINET were treated as true positive callings, and the rest were negative callings. As expected, the AUC decreased with increasing frequency cutoff (**Figure 1A, blue line**). At the frequency cutoff of 0.2, AUC reached 71.1%, but at the frequency cutoff of 0.95, the AUC lowered to 57.1%. This was consistent with the trend of total true positive callings, which declined dramatically with a high frequency cutoff **(Figure 1A, red line)**. Obviously, at lower frequency cutoff, more positives were selected and less negatives were filled in. This resulted in higher AUC, but it contained higher noise because more false positives were also added to the selection. Therefore, AUC may not be a good measurement to evaluate the accuracy of true positive calling.

**Figure 1.**
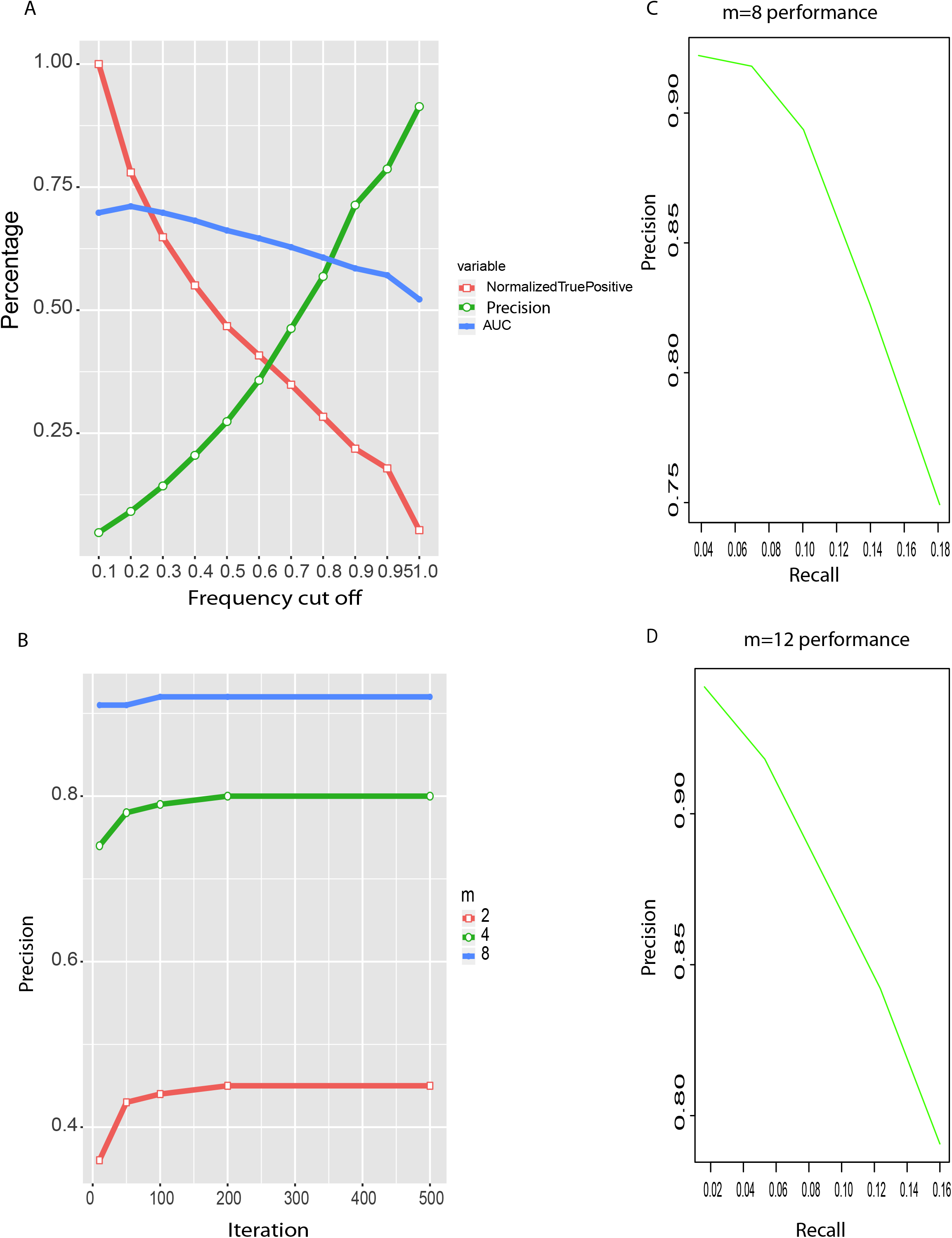
FINET parameter optimization and performance. A, Frequency cutoff optimization. Frequency cutoff from 0.1 to 1.0 vs AUC, precision and normalized true positive calling (true positive callings at each cutoff/max(true positive callings at each cutoff)). This data resulted from FINET running on network1 at dream5 with following settings, m=4, n=500, alpha=0.5 (see github software website for details). B, comparisons of precision of resampling m subgroups (frequency cutoff>0.95). C-D, the overall performance of FINET when m =8 (C) and 12 (D).

Here, we used precision (true positives/total true positive callings) to measure accuracy. During variable selection, we normally select too many variables, unsure of which one is true. In the network inference, it is more meaningful to have a higher precision than to call more true positives including noise. For example, one algorithm made 1000 true positives callings, but the real true positives were 200 (precision: 20%). Another algorithm only called 200 true positives, and 160 were real true (precision: 80%). The second algorithm missed 40 true positive (200-160), but it contained less noise than the first one, so we certainly prefer the second algorithm. In fact, some interactions in biology may not be relevant, and ignoring some interactions might make the network clear. Many biological experiments are normally conducted to prove one true gene interaction. It is valuable to obtain real true positives from computational biology. Adding false positives to get the high AUC would jeopardize the scientific value of findings. Therefore, the precision has more advantage than AUC.

The precision increased positively with frequency cutoff (**Figure 1A green line**). When frequency cutoff at 0.95, the precision reached 80% at resampling m=4. A higher frequency cutoff directly correlated to a higher precision and inversely related to the error ratio. These results fit the theory above very well. In contrast, more than 90% of true positive callings were false positives at cutoff =0.1, indicating most selections (>90%) as false without stability-selection resampling step. Therefore, the high frequency cutoff (e.g. 0.95) reduces false positive callings and makes selection stable, and stability-selection resampling is necessary.

### 3.2. Resampling m subgroups

Resampling is the key technique to improve the precision in FINET, which allows resampling m subgroups. We plotted the precision for each m (m=2,4,8) and evaluated the effect of m on the precision. When m=2, the maximum of precision only reached 45% at n=200 iterations and still kept a lot of noise, although m=2 was proposed and adopted in most current software(Meinshausen and Bühlmann, 2010; Marbach et al., 2012). This suggested that the biology is more complex than the statistic theory stated.

To solve the high noise problem, FINET increases m value as described above. FINET reaches 80% and 92% for m=4 and 8 respectively **(Figure 1B)**. In addition, when m=8, the precision reached 91% with n=10 iterations, and only slightly increased to 92% at n=100. Precision became stable at n=200. Therefore, increasing iteration n value to a big number like 10,000 as suggested in most software might not help a lot.

To appreciate the overall improvement from FINET, we plotted its precision against recall for m =8 and m =12 (**Figure 1C, 1D**). When m=8 and frequency cutoff with 0.99, 0.95, and 0,9, the precision of FINET reaches respectively 92.2%, 91.8%, and 89.4% with recall 0.04, 0.07, 0.1 (**Figure 1C**). Increasing m to 12 improves precision to 94.2%, 93.6%, 91.8% respectively for frequency cutoff of 0.99, 0.95 and 0.9, with recall 0.02, 0.04, 0.05 (**Figure 1D**). This suggested that the best way to improve accuracy is to increase sample size to allow big m value (e.g. m >=8).

## 4. Implementation, speed and usage

The whole software is implemented with Julia. From julia 0.4 to its latest version, we believe the multiple process as the stable module for parallel computations in Julia, although other approaches have been introduced. Therefore, FINET still uses multiple process modules for parallel computations. Running multiple processes requires big memory for large quantities of data. This issue is solved by using shared arrays across the processes to reduce the memory consumption in FINET.

Julia’s speed is arguably comparable to C/C++, and it runs much faster than R, Python and MATLAB, which are widely used in network inference software. For example, comparing the same Fortran code of elastic-net model, glmnet, running respectively in R and Julia for a random matrix 10000*100, Julia and R took respectively 0.7541 and 1.166 seconds to complete a single cross-validation fit. In addition, the loop over n iterations takes much longer times in R than does Julia. More importantly, it is impractical to run a data sample greater than a 20GB data matrix with R. Simply loading the 20GB data into memory (assuming memory available) takes hours for R. One can only imagine how many weeks or months it would take to select variables from a big matrix by applying algorithms with looping over n iterations in R. FINET, however, makes it possible and can complete it within hours or days in a normal computer cluster.

Using FINET is fast, accurate, easy, and accessible. FINET completes all processes with one simple command line, with input data and output file names as required, and other arguments as optional and default. The input data is a normalized matrix with each column as a gene and rows as observations (see the github web for details). Anyone with or without a computer science background can easily complete the command line.

Although developed under Linux environment, FINET should perform well in any operating system with Julia installation, including microsoftware window and apple machintosh.

